# Early urinary candidate biomarker discovery in a rat thioacetamide-induced liver fibrosis model

**DOI:** 10.1101/125120

**Authors:** Fanshuang Zhang, Yanying Ni, Yuan Yuan, Wei Yin, Youhe Gao

## Abstract

Biomarker is the change associated with the disease. Blood is relatively stable because of the homeostatic mechanisms of the body. However, urine accumulates changes of the body, which makes it a better early biomarker source. Liver fibrosis, which results from the deposition of extracellular matrix (ECM) components, is a reversible pathological condition, whereas cirrhosis, the end-stage of liver fibrosis, is irreversible. Consequently, noninvasive early biomarkers for fibrosis are desperately needed. In this study, differential urinary proteins were identified in the thioacetamide (TAA) liver fibrosis rat model using tandem mass tagging and two-dimensional liquid chromatography tandem mass spectrometry (2DLC-MS/MS). A total of 766 urinary proteins were identified, 143 and 118 of which were significantly changed in the TAA 1-week and 3-week groups, respectively. Multiple reaction monitoring (MRM)-targeted proteomics was used to further validate the abundant differentially expressed proteins in the TAA 1-week, 3-week, 6-week and 8-week groups. A total of 40 urinary proteins were statistically significant (fold change >2 and p<0.05), 15 of which had been previously reported as biomarkers of liver fibrosis, cirrhosis or other related diseases and 10 of which had been reported to be associated with the pathology and mechanism of liver fibrosis. These differential proteins were detected in urine before the alanine aminotransferase (ALT) and aspartate transaminase (AST) changes in the serum and before fibrosis was observed upon hematoxylin and eosin (HE) and Masson’s staining.

## Introduction

Liver fibrosis is a state of liver injury in which excessive extracellular matrix (ECM) components, especially collagen, are deposited and disturb normal liver functions, which results in many chronic liver diseases ^[1, 2]^. During the process of chronic liver injury, fibrotic scar tissue is gradually formed due to excessive deposition, hepatic architecture is distorted, and nodules of regenerating hepatocytes are ultimately generated, which result in cirrhosis ^[3, 4]^. Liver fibrosis is a reversible pathological condition, whereas cirrhosis, the end-stage of liver fibrosis, is irreversible ^[5, 6]^. Cirrhosis can affect the risk of developing primary liver cancer or hepatocellular carcinoma (HCC) ^[7]^. Consequently, noninvasive biomarkers that have adequate specificity and sensitivity and respond quickly to changes in the fibrogenic process are desperately needed ^[3]^.

Liver fibrosis and cirrhosis are commonly studied using rat models ^[8]^. Among the liver fibrosis animal models, which are induced by bile duct ligation (BDL) ^[9]^, ethanol ^[10]^, carbon tetrachloride (CCl_4_) ^[11]^ or thioacetamide (TAA) ^[12]^, the intraperitoneally TAA-injected rat model is widely used to induce liver fibrosis and cirrhosis and consistently produces liver fibrosis and cirrhosis in rats with a histopathology that is more similar to that of human liver fibrosis and cirrhosis ^[13-15]^. In 1948, TAA was first reported as a hepatotoxic agent ^[16]^. Chronic TAA application was shown to lead to centrilobular necrosis and substantial liver fibrosis and cirrhosis in rats ^[17-19]^. The advantages of the intraperitoneally TAA-injected rat model include its high specificity for the liver and a large window of time before liver fibrosis and cirrhosis ^[20, 21]^.

Biomarker is the change associated with the disease. Blood is relatively stable because of the homeostatic mechanisms of the body. However, urine accumulates changes of the body, which makes it a better early biomarker source ^[22, 23]^. Urinary proteomics has become increasingly important in studies of quantitative changes in proteins resulting from changes in disease states ^[24-28]^; moreover, numerous urinary protein biomarkers have been reported in different diseases ^[29-32]^. These findings suggest that urinary proteins may serve as non-invasive biomarkers for diseases.

However, the composition of urine is influenced by multiple pathological and physiological factors, such as gender, age, and medicine ^[33]^. To minimize the affecting factors, animal models can be used for urinary biomarker discovery of liver fibrosis ^[34]^.

In the present study, a rat model of TAA-induced liver fibrosis and cirrhosis was used to identify urinary protein biomarkers related to the developmental process (1 week, 3 weeks, 6 weeks and 8 weeks) of liver fibrosis using urinary proteomic profiling.

## Materials and methods

### Experimental animals

Male Sprague-Dawley rats 6–8 weeks old and weighing 180–200 g were purchased from the Institute of Laboratory Animal Science, Chinese Academy of Medical Science & Peking Union Medical College. All animal protocols governing the experiments in this study were approved by the Institute of Basic Medical Sciences Animal Ethics Committee, Peking Union Medical College (Approved ID: ACUC-A02-2014-008). All animals were maintained with a standard laboratory diet under controlled indoor temperature (22 ± 1°C) and humidity (65 ∼ 70%) and with a 12-h light-dark cycle. The study was performed according to the guidelines developed by the Institutional Animal Care and Use Committee of Peking Union Medical College. All efforts were made to minimize suffering.

### Liver fibrosis and cirrhosis model

Rats were divided randomly into two groups. The TAA group (n = 22) was injected intraperitoneally with 200 mg/kg TAA thrice weekly for eight weeks to establish the liver cirrhosis model. The control group (n = 20) was injected intraperitoneally with equivalent volumes of saline. The body weights of the rats were recorded weekly. The urine samples from ten rats in each group were individually collected in metabolic cages at weeks 1, 3, 6 and 8. During urine collection, in order to avoid contamination, all rats were given free access to water without food. The urinary protein and creatinine concentrations were assayed spectrophotometrically at the Peking Union Medical College Hospital.

### Serum biochemical parameters

After the urine samples were collected, five rats from each group were anesthetized with 2% pelltobarbitalum natricum (40 mg/kg body weight), and 2–3 mL of blood was withdrawn through the abdominal aorta in a heparinized tube and centrifuged at 2000×g for 20 min at 4°C to obtain serum. The aspartate aminotransferase (AST), alanine aminotransferase (ALT), alkaline phosphatase (ALP), albumin, total protein and creatinine concentrations were assayed spectrophotometrically at the Peking Union Medical College Hospital.

### Histopathology

The liver samples were washed in ice-cold saline, blotted on filter paper, weighed, fixed in 10% neutral-buffered formalin, and underwent histopathology. To determine whether the kidneys were injured, kidney samples were also fixed in 10% neutral-buffered formalin for histopathology. The formalin-fixed tissues were embedded in paraffin, sectioned at 3–5 mm and stained with hematoxylin and eosin (HE) to reveal histopathological lesions. Liver fibrosis was evaluated by Masson’s trichrome staining ^[35]^.

### Immunohistochemistry

The paraffin-embedded sections from individual samples were permeabilized with 0.2% Triton and blocked with 5% bovine serum albumin (BSA) in 0.1 M phosphate-buffered saline (PBS) for 30 min to reduce nonspecific binding. Then, the samples were incubated with primary antibodies against Vimentin (15200, ab9547, Abcam, Cambridge, MA, USA) followed by a biotinylated secondary antibody (PV-9000, Beijing ZSGB-Bio, Beijing, China). The samples were incubated with the IgG K-light chain (15200, M0809-1, Hangzhou HuaAn Biotechnology Company, Hangzhou, China) at a similar concentration as the primary antibody controls, followed by the same biotinylated secondary antibody (PV-9000, Beijing ZSGB-Bio, Beijing, China). Immunoreactivity was visualized with diaminobenzidine (DAB), and brown staining was considered a positive result.

### Urinary protein sample preparation

Urine was centrifuged at 2,000×g for 15 min at 4°C. After the cell debris was removed, the supernatant was centrifuged at 12,000×g for 15 min at 4°C. Three volumes of ethanol were added after removing the pellets and precipitated at 4°C. After the supernatant was removed, the pellets were re-suspended with lysis buffer (8 M urea, 2 M thiourea, 25 mM dithiothreitol (DTT) and 50 mM Tris) ^[36]^. Protein concentrations were measured using the Bradford method. Proteins were digested with trypsin (Trypsin Gold, Mass Spec Grade, Promega, Fitchburg, Wisconsin, USA) using filter-aided sample preparation methods ^[37]^. Briefly, after proteins were loaded onto a 10-kDa filter unit (Pall, Port Washington, New York, USA), UA buffer (8 M urea in 0.1 M Tris-HCl, pH 8.5) and NH_4_HCO_3_ (25 mM) were added successively, and the tube was centrifuged at 14,000×g for 20 min at 18°C. Proteins were denatured by incubation with 20 mM DTT at 50°C for 1 h and then alkylated with 50 mM iodoacetamide (IAA) for 45 min in the dark. After the samples were centrifuged with UA twice and NH_4_HCO_3_ four times, the proteins were redissolved in NH_4_HCO_3_ and digested with trypsin (1:50) at 37°C overnight. The tryptic peptides were desalted using Oasis HLB cartridges (Waters, Milford, Massachusetts, USA). Finally, the desalted peptides were dried by vacuum evaporation (Thermo Fisher Scientific, Bremen, Germany).

### Peptide tandem mass tag (TMT) labeling

The peptides were solubilized in 100 mM tetraethylammonium bromide (TEAB) and labeled with the 6-plex Tandem Mass Tag Label Reagents provided by Thermo Fisher Scientific (Pierce, Rockford, IL, USA), which were equilibrated to room temperature immediately before use. Then, 41 μL anhydrous acetonitrile was added to each tube, and the reagent was allowed to dissolve for 5 min with occasional vortexing. The samples were briefly centrifuged to gather the solution, and 20 μL of the TMT Label Reagent was added. The reaction was then incubated for 2 h at room temperature. After the peptides were labeled with isobaric tags, they were mixed at a 1:1:1:1:1:1 ratio based on the amount of total peptide, which was determined by running an equal volume proportion of labeled samples using liquid chromatography coupled with tandem mass spectrometry (LC-MS/MS) and comparing the total signal intensities of all peptides. Finally, the control group (n = 5), TAA 1-week group (n = 5), TAA 3-week group (n = 5) and one pooled sample (mixture of 15 samples) were analyzed by two-dimensional LC-MS/MS (2DLC-MS/MS).

### HPLC separation

The TMT-labeled samples were fractionated using a high-pH reversed-phase liquid chromatography (RPLC) column from Waters (4.6 mm × 250 mm, Xbridge C18, 3 μm) and loaded onto the column in buffer A1 (H2O, pH=10). The elution gradient was 5–;25% buffer B1 (90% ACN, pH=10; flow rate=1 mL/min) for 60 min. The eluted peptides were collected at one fraction per minute. The 60 dried fractions were re-suspended in 0.1% formic acid and pooled into 30 samples by combining fractions 1 and 31, 2 and 32, etc. The odd-numbered fractions were chosen for further analysis. A total of 45 fractions from urinary peptide mixtures were analyzed by LC-MS/MS.

### LC-MS/MS analysis

Each fraction was analyzed with a reverse-phase-C18 self-packed capillary LC column (75 μm × 100 mm). The eluted gradient was 5%–30% buffer B2 (0.1% formic acid, 99.9% ACN; flow rate=0.3 μL/min) for 40 min. A Triple TOF 5600 mass spectrometer was used to analyze the eluted peptides from LC, and each fraction was run twice. The MS data were acquired using the high-sensitivity mode with the following parameters: 30 data-dependent MS/MS scans per full scan, full scans acquired at a resolution of 40,000, MS/MS scans at a resolution of 20,000, rolling collision energy, charge state screening (including precursors with a charge state of +2 to +4), dynamic exclusion (exclusion duration 15 s), an MS/MS scan range of 100-1800 m/z, and a scan time of 100 ms.

### Data analysis

The MS/MS data were subjected to Mascot software (version 2.4.1, Matrix Science, London, UK) analysis, and proteins were identified by comparing the peptide spectral matches against the Swissprot_2014_07 databases (taxonomy: Rattus, containing 7,906 sequences). Trypsin was selected as the digestion enzyme with up to two missed cleavage sites allowed, and carbamidomethylation (57.02146) on a cysteine was defined as a fixed modification. The precursor ion mass tolerance and the fragment ion mass tolerance were 0.05 Da. Protein identification of the Mascot results was validated by using Scaffold Proteome Software (version Scaffold_4.3.3, Proteome Software Inc., Portland, OR). Peptide identification was accepted at a false discovery rate (FDR) of less than 1.0% at the protein level and contained at least 2 unique peptides. Scaffold Q+ was used for the quantification of Label-Based Quantification (TMT, iTRAQ, SILAC, etc.) peptides and proteins. Reporter ion intensities acquired in each channel were normalized by the sum of all reporter ion intensities of the corresponding channel. Normalized reporter ion intensities were used to calculate the relative protein abundance. Then, the protein ratios were quantified by the median of the transformed reporter ion intensity ratios ^[38, 39]^.

### Multiple reaction monitoring (MRM) confirmation

Multiple reaction monitoring (MRM) was employed to analyze the resulting TMT-labeled mass spectrometer data of the significantly changed proteins. Data derived from a spectral library of the urinary proteomics generated by conventional LC-MS/MS using HCD collision were imported into the Skyline software (version 1.1). Skyline was applied to select the most intense transitions for the targeted peptides ^[40, 41]^. The b and y ions of fragments exceeding the m/z ratio of doubly and triply charged peptide precursors were considered. A maximum of five transitions per peptide were traced on a QTRAP 6500 mass spectrometer (AB Sciex). The ideal peptides from the target list for building MRM were further optimized using the following criteria: 1) the peptide had no missed cleavage site with trypsin, 2) the peptide was unique to one protein, and 3) the peptide did not contain asparagine, glutamine, methionine. We used the urine samples from the control 1-week group (n = 5), control 3-week group (n = 5), control 6-week group (n = 5), control 8-week group (n = 5), TAA 1-week group (n = 5), TAA 3-week group (n = 5), TAA 6- week group (n = 5) and TAA 8-week group (n = 5) for the MRM confirmation. Approximately 200 μg of each urinary protein sample was digested by trypsin through centrifugation in a 10-kDa filter unit (Pall, Port Washington, New York, USA). All tryptic peptides were loaded onto a self-packed C18 RP capillary column (100 mm × 0.075 mm, 3 μm) with buffer A (0.1% formic acid). The peptides were eluted with 5–30% buffer B (0.1% formic acid, 99.9% ACN; flow rate=300 nL/min) for 60 min. Each sample was run in triplicate. All of the MS data were loaded into Skyline for further visualization, transition detection, and abundance calculations.

### Statistical analysis

All statistical analyses were performed with the Statistical Package for Social Studies software (SPSS, version 16, IBM), and a p-value less than 0.05 was considered statistically significant. Comparisons between independent groups were conducted using one-way ANOVA followed by post hoc analysis with the least significant difference (LSD) test or Dunnett’s T3 test.

## Results

### Body weight

Rats from the control group exhibited a normal weight gain and followed by a normal growth pattern. However, after one week of TAA injection, the rats of the TAA group suffered growth retardation and had a significantly lower weight than the group (p < 0.05). The continuous change in body weight in the control 8-week group (n = 5) and TAA 8-week group (n = 7) is shown in Figure 1A, and the liver index of the sacrificed rats (liver weight/body weight) in the control 1-week group (n = 5), control 3-week group (n = 5), control 6-week group (n = 5), control 8-week group (n = 5), TAA 1-week group (n = 5), TAA 3-week group (n = 5), TAA 6-week group (n = 5), and TAA 8-week group (n = 5) is shown in Figure 1B.

**Figure 1.**
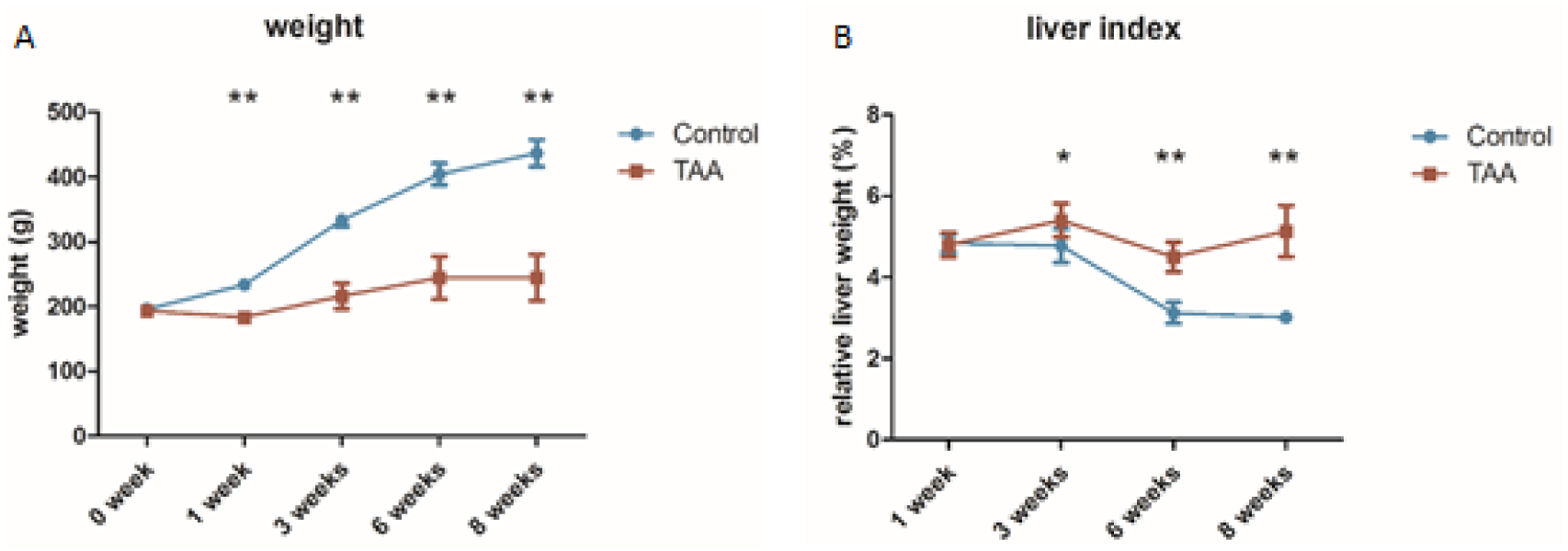
A, The continuous change in body weight in the control 8-week group (n=5) and TAA 8-week group (n=7). B, The liver index of the sacrificed rats (liver weight/body weight) in the control 1-week group (n=5), control 3-week group (n=5), control 6-week group (n=5), control 8-week group (n=5), TAA 1-week group (n=5), TAA 3-week group (n=5), TAA 6-week group (n=5), and TAA 8-week group (n=5). p<0.05 (*), p<0.01 (**).

### Serum biochemical parameters

The serum levels of specific liver function biomarkers were assayed to determine the liver function of each rat. There was no difference in serum biochemical parameters in the TAA 1-week group and 3-week group compared with those in the respective control groups. However, the TAA 6-week and TAA 8-week groups showed a significant increase in the levels of liver function biomarkers ALT and aspartate transaminase (AST) and a significant decrease in the levels of total protein (TP) and albumin (ALB) compared to those in the respective control groups, indicating hepatocyte damage (Figure 2).

**Figure 2.**
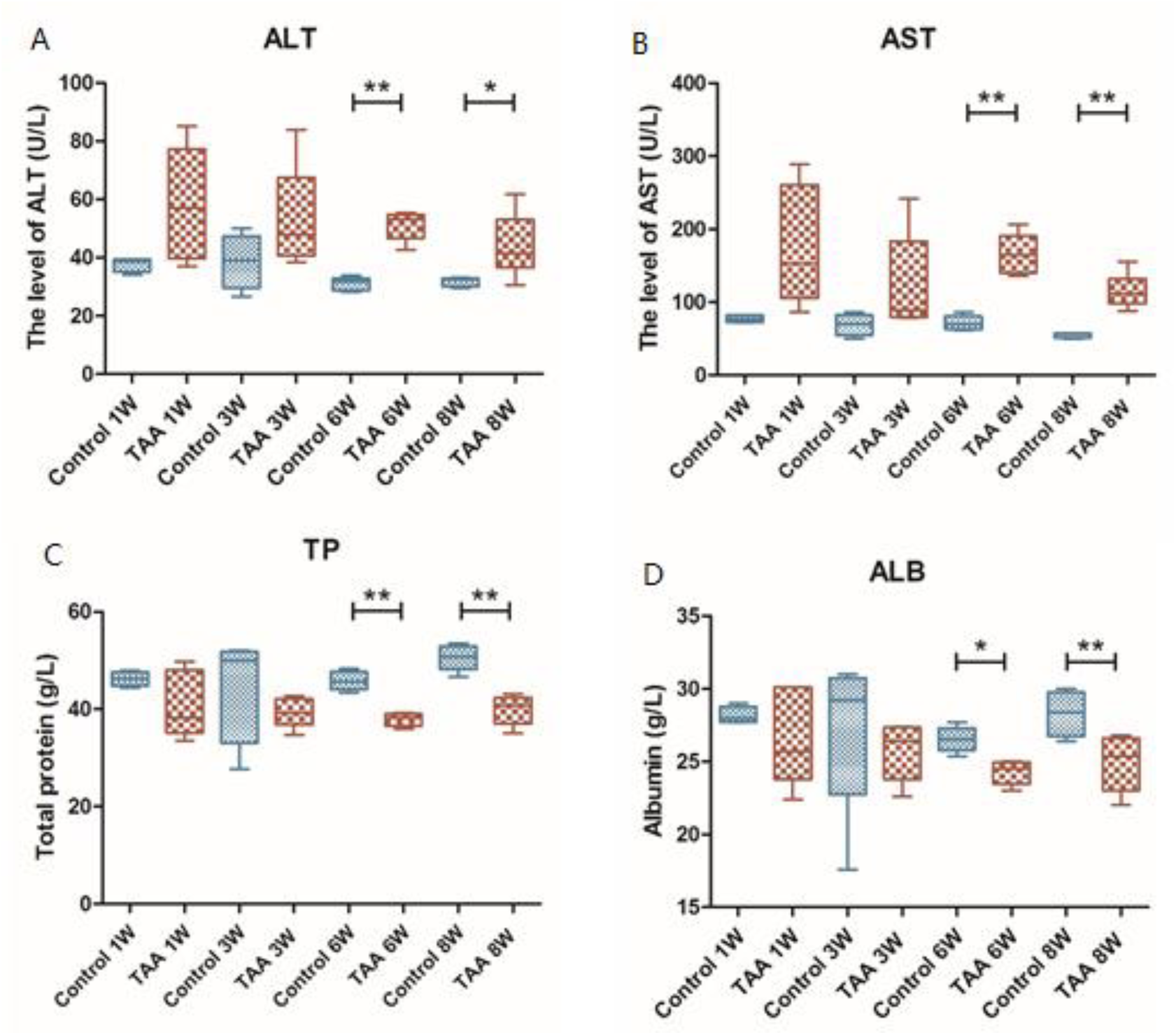
Serum Biochemical Parameters. A, The level of ALT. B, The level of AST. C, The level of TP. D, The amount of ALB. p<0.05 (*), p<0.01 (**).

### Histopathology

The HE staining and Masson’s trichrome staining of liver tissue obtained from the control group and TAA groups are shown in Figure 3. No histological lesions were seen in the control group. In the TAA 1-week group and 3-week group, the liver tissue showed an almost normal lobular architecture with distinct hepatic cells, a central vein, and sinusoidal spaces. However, in the TAA 6-week group, some inflammatory cells, collagen deposition, and hydropic degeneration of endothelial cells and hepatocytes were observed mainly in the centrilobular areas forming thin fibrous septa around the central veins, whereas the hepatic lobules were almost well-arranged. In the TAA 8-week group, fibrous bridges were completely formed between the central veins and the central veins to the portal areas, thus separating the liver parenchyma into a typical pseudo-lobule. The fibrous bridges (the fibrotic lesions) became thicker, resulting in complete cirrhosis. Collagen accumulation and cirrhotic nodules were shown by using Masson’s trichrome staining. The HE staining of the renal tissue obtained from the TAA 8-week group showed that neither the tubules nor the glomeruli were damaged, suggesting that the differences between the TAA group and control group were caused by cirrhosis, not kidney damage (Figure 4).

**Figure 3.**
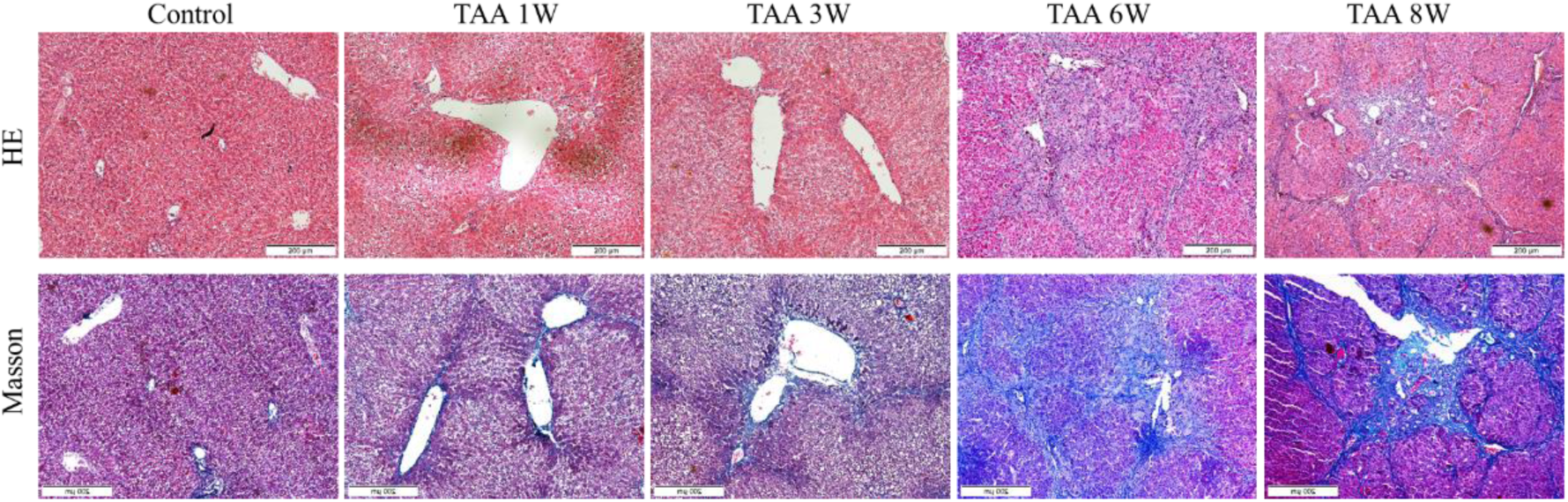
Pathological morphologies of the liver in the control and TAA groups. A, HE staining of rat livers in the control group and TAA group at 1, 3, 6 and 8 weeks of TAA injection. B, Masson’s trichrome staining of rat livers in the control group and TAA group at 1, 3, 6 and 8 weeks of TAA injection

**Figure 4.**
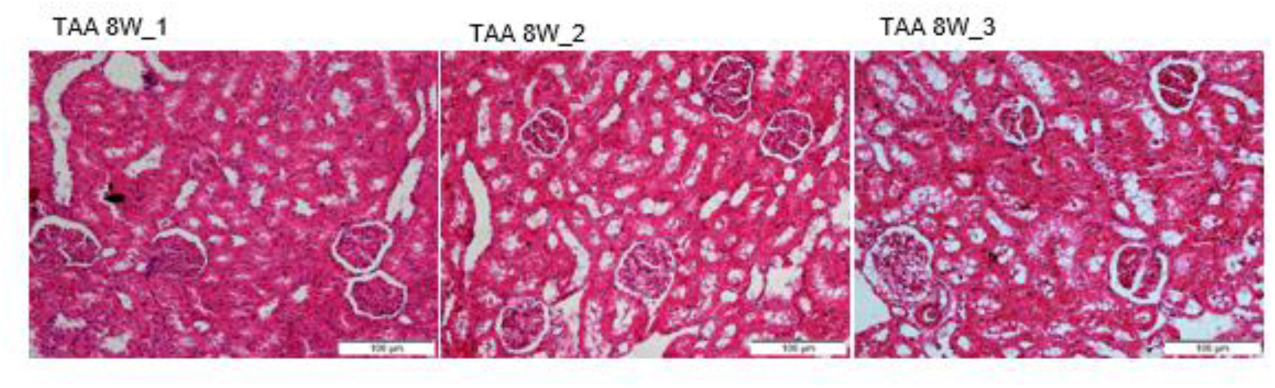
Pathological morphologies of the kidney in the control and TAA groups. A, HE staining of rat kidneys in the TAA 8-week group.

### Urinary proteome changes identified by LC-MS/MS

In the present study, the urine samples of 15 rats in the control group (n=5), TAA 1-week group (n=5) and TAA 3-week group (n=5) were digested by trypsin and labeled with the 6-plex TMT reagent; the pooled urinary sample (mixture of the 15 samples) was used as a control and analyzed by 2DLC-MS/MS twice. For the total difference in the spectra between the TAA group and the control group, spectral counting was used to perform a semi-quantitative analysis ^[35, 42]^. The abundance of each protein in a sample was estimated by the mean spectral count of two replicates.

At the protein level, a total number of 766 protein groups was identified in the urinary proteome at an FDR < 1%, including at least 2 unique peptides, and 467 proteins could be quantified in all 15 urine samples in the technical replicates. Relative to the control group, 143 and 118 significantly changed urinary proteins were identified in the TAA 1-week and 3-week groups, respectively (Tables S1, S2), with 90 proteins overlapping between these two sets of significantly changed proteins (Figure 5). To be conservative, all of the significantly changed proteins met the following criteria: 1) the proteins had at least two unique peptides, 2) the variation trend of the proteins in all five animals of each group was consistent, and 3) the fold change was more than 2.

**Figure 5.**
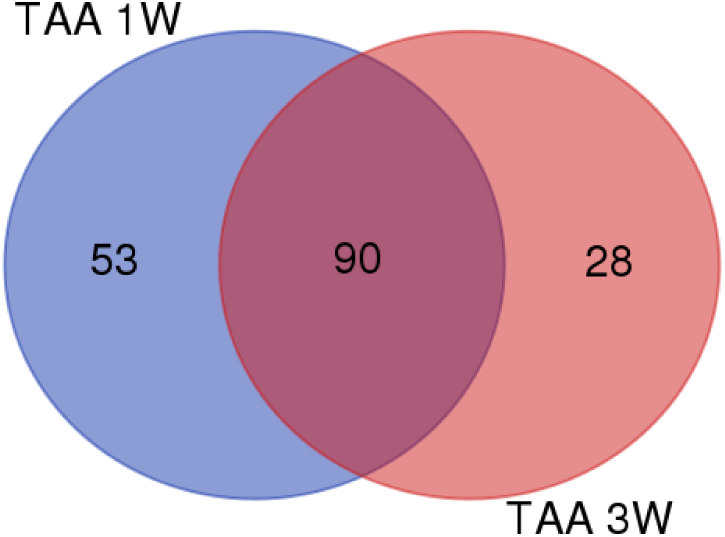
Significantly changed urinary proteins in the TAA 1 week and 3-week groups.

### Clustering of proteins

A hierarchical clustering was performed by using the average linkage method. As shown in Figure 6, all 467 proteins were clustered into 3 clusters, which corresponded to the control group, TAA 1-week group and TAA 3-week group (Figure 6), and all technical replicates within one sample were clustered together, demonstrating that the technical variation was smaller than the inter-individual variation. Moreover, all 5 samples from the same group could be clustered together. These results indicated that the intra-group technical variation was smaller than the inter-group biological variation.

**Figure 6.**
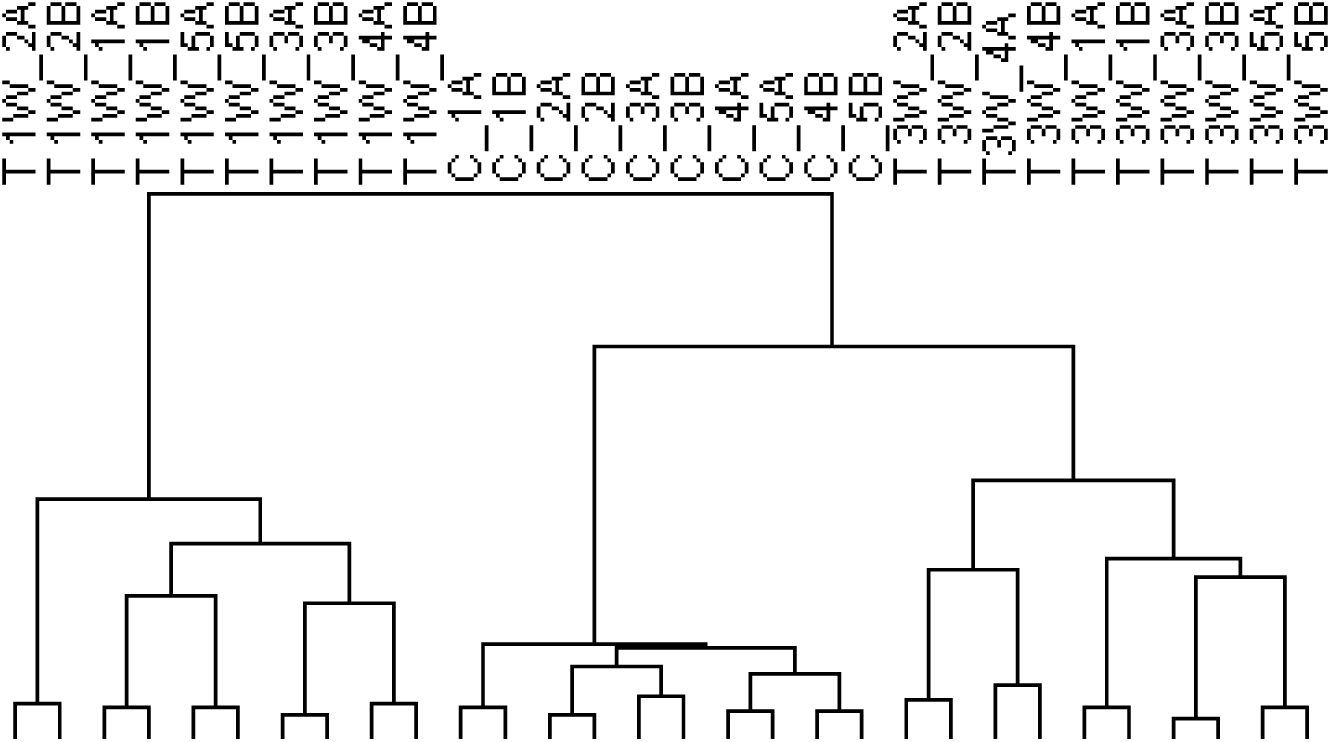
Analysis of the protein clustering identified by the TMT method.

### Multiple reaction monitoring (MRM) analysis

For further validation of the differentially abundant proteins, we analyzed the individual TAA group and control group urine samples using the MRM targeted proteomics method. From the significantly changed urinary proteins, 63 proteins were chosen for validation, and the data were analyzed by the Skyline software ^[41]^. After the ideal peptides were further optimized for creating the MRM analysis, 57 proteins (47 increased proteins and 10 decreased proteins) were finally used for validation using MRM targeted proteomics. The technical reproducibility of each individual MRM assay was assessed, and the CV values are shown in Figure 7. Overall, 51 of the 57 investigated proteins (47 increased proteins and 4 decreased proteins) exhibited the same average trend in the differential abundance of the proteins observed by both high-throughput analysis and the MRM method. Moreover, 40 proteins were statistically significant (p<0.05) with a fold change greater than 2 according to their abundance among groups, strongly supporting their potential clinical relevance in liver fibrosis (Table 1). Figure 8 shows several of these proteins.

**Figure 7.**
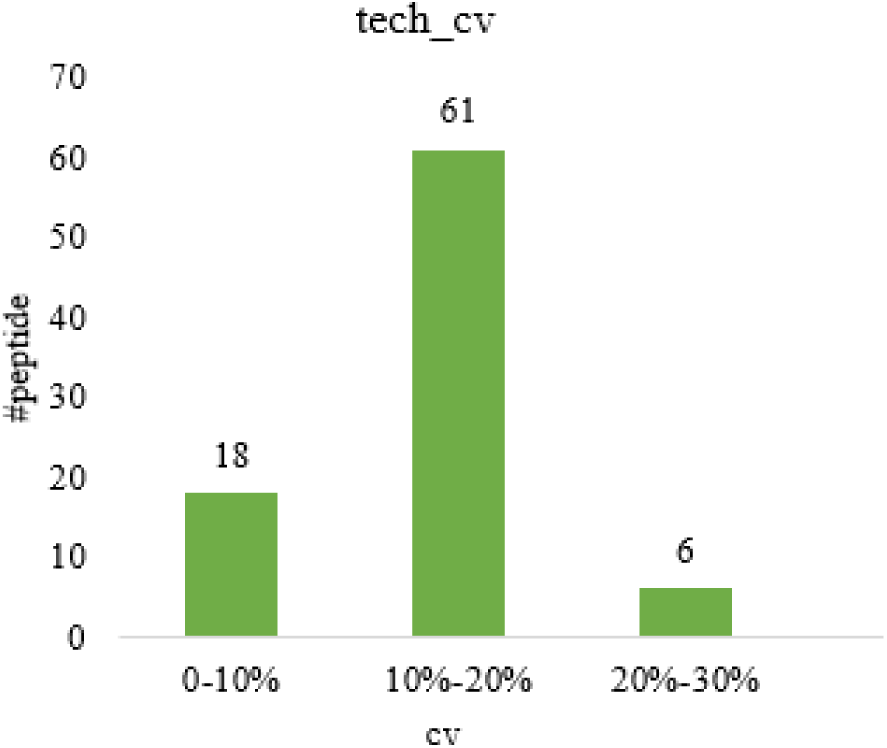
The CV value of the technical reproducibility of each individual MRM assay.

**Table 1.**
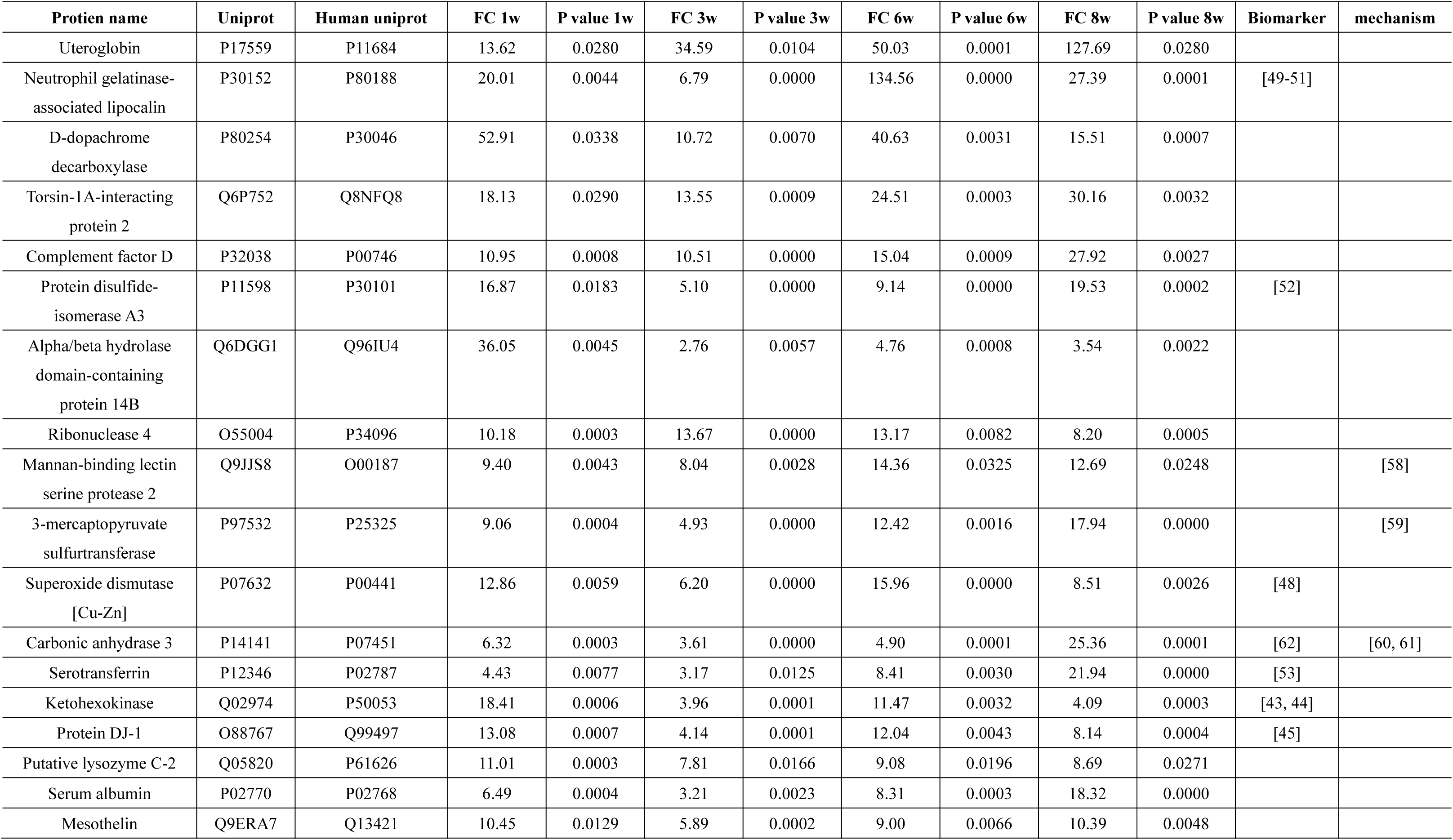
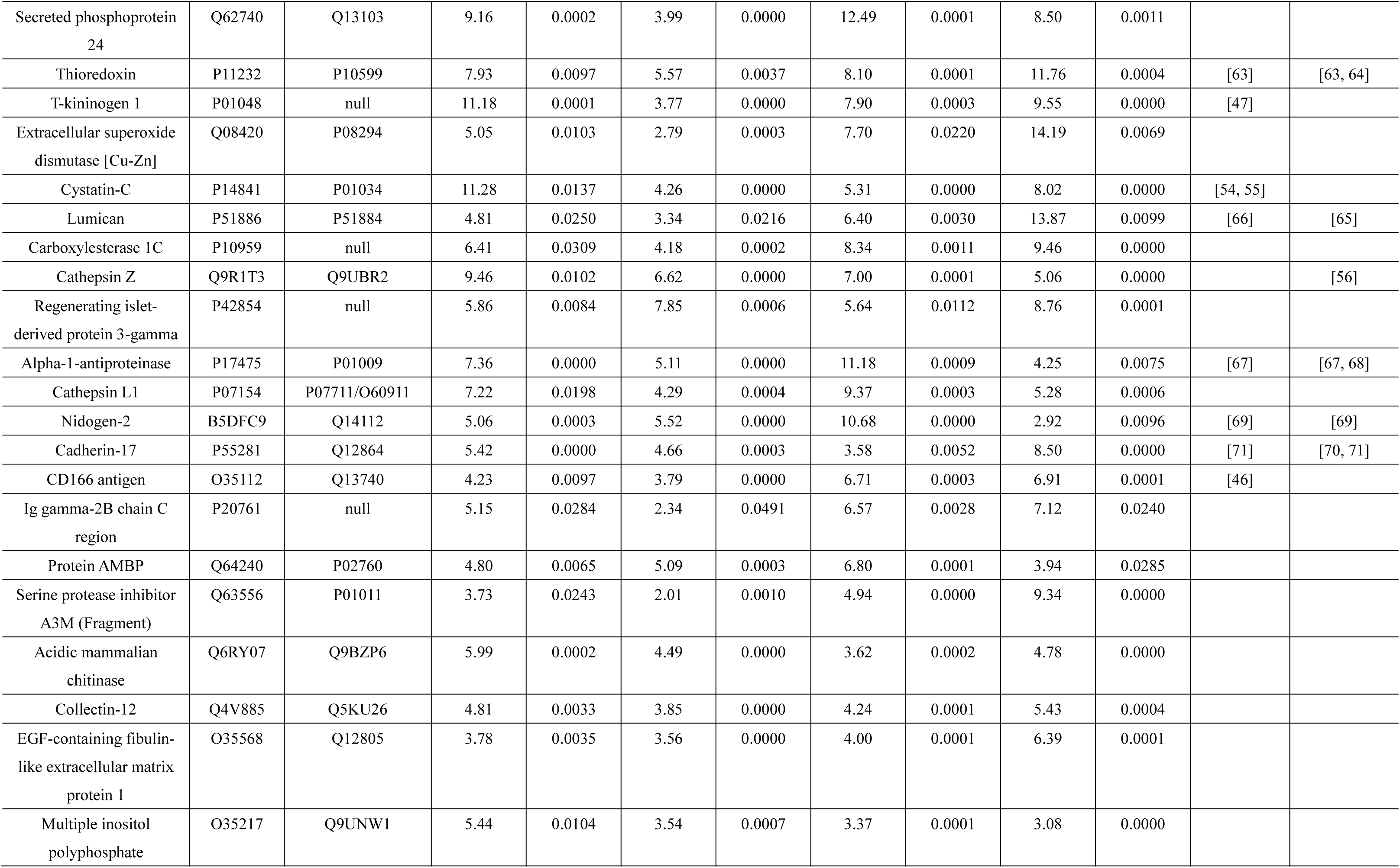
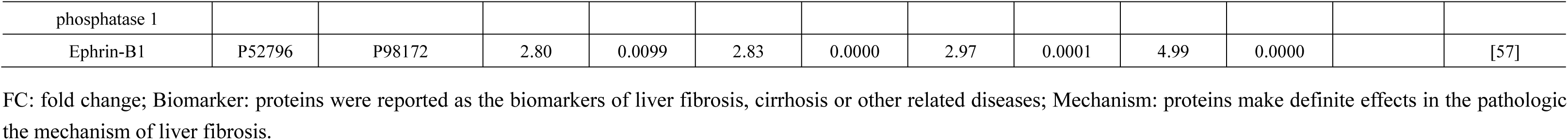
Details of changed urinary proteins identified in the TMT method.

**Figure 8.**
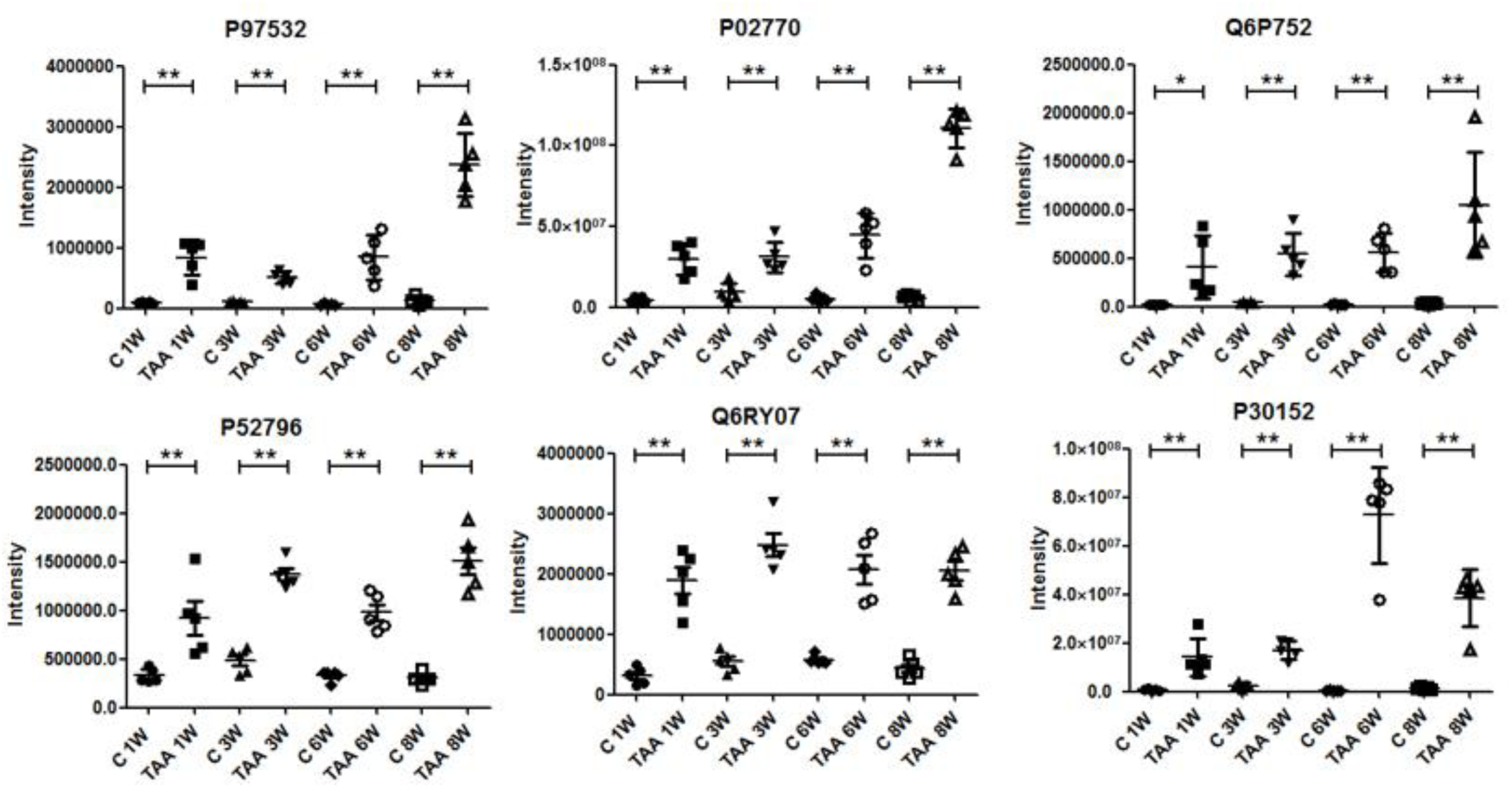
The intensity of several significantly changed urinary proteins validated by MRM. p<0.05 (*), p<0.01 (**).

Among the 51 MRM-verified significantly changed proteins, some had previously been reported as biomarkers of liver fibrosis, cirrhosis or other related diseases. The expression level of ketohexokinase is clearly up-regulated in mice with Con A-induced hepatitis ^[43]^ and is one of the potential hepatocarcinogenic biomarkers ^[44]^. Protein DJ-1expression is significantly upregulated in hepatocellular carcinoma (HCC) and its expression level correlates with clinicopathological variables and prognosis of HCC patients, which suggests that DJ-1 maybe a candidate prognostic biomarker of HCC ^[45]^. CD166 antigen is a novel tumor marker of HCC ^[46]^. T-kininogen 1 is a new potential serum biomarker for inflammatory hepatic lesions ^[47]^. Extracellular superoxide dismutase activities in liver homogenates is significantly decreased in the CCl_4_-treated liver fibrosis rat model ^[48]^. Neutrophil gelatinase-associated lipocalin is a urine biomarker of acute-on-chronic liver failure and cirrhosis ^[49-51]^. Protein disulfide-isomerase A3 is a biomarker for liver fibrosis caused by CCl_4_ in rats ^[52]^. Serotransferrin is a biomarker that is decreased in liver fibrosis serum ^[53]^. Finally, cystatin C has been identified as a non-invasive serum marker of liver fibrosis ^[54, 55]^.

Some of the significantly changed proteins identified here have been reported to relate to the development of liver fibrosis or other related diseases. Overexpression of cathepsin Z contributes to tumor metastasis by inducing epithelial-mesenchymal transition in HCC ^[56]^. Ephrin-B1 may be involved in in vivo tumor progression by promoting neovascularization in HCC ^[57]^. Mannan binding lectin-associated serine protease activates human hepatic stellate cells, which is activated in the pathogenesis of liver fibrosis ^[58]^. 3-mercaptopyruvate sulfurtransferase (MST) is expressed in the liver and regulates liver functions via H_2_S production, and malfunction of hepatic H_2_S metabolism may be involved in many liver diseases, such as liver fibrosis and liver cirrhosis ^[59]^.

What’s more, several proteins have been previously reported to be associated with the pathology and mechanism of liver fibrosis, cirrhosis and other related diseases as well as the biomarkers of these diseases. Carbonic anhydrase 3 promotes transformation and invasion capability in hepatoma cells through FAK signaling pathway^[60]^ and is a major participant in the liver response to oxidative stress ^[61]^. Carbonic anhydrase 3 is also a biomarker of liver injury ^[62]^. Thioredoxin has a potential to attenuate liver fibrosis via suppressing oxidative stress and inhibiting proliferation of stellate cells ^[63]^ and plays important roles in the pathophysiology of liver diseases ^[64]^. Overexpression of thioredoxin has been reported to prevent TAA-induced hepatic fibrosis in mice ^[63]^. Lumican is a prerequisite for liver fibrosis ^[65]^ and the altered expression of lumican has been associated with liver fibrosis ^[66]^. Alpha-1-antiproteinase (A1AT) is the most abundant liver-derived glycoprotein in plasma. The deposition of excessive A1AT in liver cells is associated with increased risk for liver cirrhosis ^[67]^. Hereditary deficiency of A1AT in plasma can cause liver disease in childhood and cirrhosis and/or hepatocellular carcinoma (HCC) in adulthood ^[68]^. The level of A1AT has been found significantly high in liver cirrhosis with hepatitis C viral infection and HCC patients but less in chronic hepatitis C than control subjects and can be used as biomarkers for monitoring the liver diseases ^[67]^. Nidogen-2 has been reported significantly decreased in HCC tissues, which is significantly correlated with tumor progression factors. The decreased expression of nidogen-2 may have a potential pathogenetic role in the development of HCC ^[69]^. Furthermore, nidogen-2 also decreases in serum of HCC and may also have potential diagnostic value for HCC ^[69]^. Cadherin-17 (CDH17), also called liver-intestine cadherin, is expressed in the liver and intestinal epithelial cells and have a role in the morphological organization of liver and intestine ^[70]^. CDH17 is an oncofetal molecule of HCC by exhibiting elevated expression during embryogenesis and carcinogenesis of the livers and has the potential applicability to be a molecular diagnosing biomarker and target for HCC ^[71]^.

There were also some differential proteins discovered in our study that have never been reported to relate to liver fibrosis, such as torsin-1A-interacting protein 2, putative lysozyme C-2, and cathepsin L1. Since these proteins were changed dramatically, they also have the potential to be early urinary biomarkers of liver fibrosis.

## Discussion

Liver fibrosis is one of the major health problems in the world ^[72]^. What is more, liver fibrosis is still reversible, but when it develops into cirrhosis, which is the irreversible end-stage of liver fibrosis, recovery is impossible ^[49, 58]^. For now, liver puncture biopsy is still the gold standard for the diagnosis of liver fibrosis; therefore, the discovery of noninvasive biomarkers is increasingly important ^[73]^. Urine is being recognized as a good source for noninvasive biomarkers because it accumulates the pathological and physiological changes of the body, which are the most fundamental properties of biomarkers ^[34]^. In this study, the TAA-induced liver fibrosis rat model was utilized to simulate the progression of liver fibrosis. This method enabled the identification of differentially expressed proteins by urinary proteomic profiling.

Here, the urinary proteomes of rats exposed to TAA for 1 week and 3 weeks were profiled using the TMT-labeled LC-MS-MS method. Several differential proteins were identified between the TAA groups and the control group. The MRM validation of the differentially expressed proteins of the TAA 1-week, 3-week, 6-week and 8-week groups and their relevant control groups further confirmed the results of our proteome analysis. Some of the identified urinary proteins showed a significant expression change (p < 0.05) even in the TAA 1-week group, which was earlier than the appearance of the changes in ALT and AST in the serum and fibrosis in the liver using HE and Masson’s staining. Importantly, these results indicated that these urinary proteins have the potential to be a better noninvasive biomarker of liver fibrosis. However, the body weight of the rats was also significantly different in the TAA 1-week group, potentially due to the toxicity of TAA or the influence of the digestive system function, and this change was not specific to liver fibrosis. By decreasing the dosage of TAA according to the body weight of the rats, thereby slowing the disease progression, we may be able to identify earlier changes in urinary proteins before the change in the body weight. In this preliminary study, only one type of liver fibrosis animal model was used for the discovery of urinary biomarkers. Further analysis of other animal models or a large number of clinical samples should also be used to verify a specific protein or a panel of proteins as clinically applicable biomarkers of liver fibrosis.

## Acknowledgments

This work was supported by the National Key Research and Development Program of China (2016YFC1306300); the National Basic Research Program of China (2013CB530850); Beijing Natural Science Foundation (7173264, 7172076) and funds from Beijing Normal University (11100704, 10300-310421102).

